# Repurposing tRNA isodecoders for non-canonical functions via tRNA cleavage

**DOI:** 10.1101/2024.09.04.611212

**Authors:** Nupur Bhatter, Vivek M. Advani, Yoshika Takenaka, Shawn M. Lyons, Yasutoshi Akiyama, Paul J. Anderson, Pavel Ivanov

**Affiliations:** Division of Rheumatology, Inflammation and Immunity, Brigham and Women’s Hospital, Boston, MA, USA; Department of Medicine, Harvard Medical School, Boston, MA, USA; Laboratory of Oncology, Pharmacy Practice and Sciences, Tohoku University Graduate School of Pharmaceutical Sciences, Sendai, Japan; Department of Biochemistry, Boston University Medical School, Boston, USA; Harvard Initiative for RNA Medicine, Boston, MA, USA

**Keywords:** tRNA, tRNA isodecoders, tRNA cleavage, protein synthesis, ribonuclease, tiRNAs, tDRs

## Abstract

Transfer RNAs (tRNAs) are the key adaptor molecules aiding protein synthesis. Hundreds of tRNA genes are found in the human genome but the biological significance of this genetic excess is still enigmatic. The tRNA repertoires are variable between tissues and cells as well as during development. Such variations can only be partially explained by the correlation to the physiological needs in protein production, e.g. by changes in the expression of tRNA isoacceptor sets (tRNAs charged with the same amino acid but bearing different anticodons). However, changes in the expression levels of individual isodecoders (tRNAs with the same anticodon) are less understood. Besides canonical functions in mRNA translation, tRNAs are implicated in non-canonical functions unrelated to protein synthesis. tRNAs are rich source of small non-protein coding RNAs called tRNA-derived RNAs (tDRs), which include tRNA-derived stress-induced RNAs (tiRNAs) formed in response to stress. Here, we show that tiRNAs derived from isodecoders different in a single nucleotide can also differ in their activities. Specifically, we show that isodecoder sets for tRNA^His-GTG^, tRNA^Gly-GCC^ and tRNA^Cys-GCA^ are cleaved by ribonucleases to yield 5’-tiRNAs showing differential activity towards mRNA reporter translation. Our data propose a model where cleavage repurposes specific tRNA isodecoders for non-canonical functions.

**Significance Statement:** The human genome encodes hundreds of transfer RNA (tRNA) genes to decode 61 codons. The basis for such genetic redundancy is unclear but the increase in the number of tRNA genes goes in concert with the complexity of an organism. While changes in the expression of isoacceptor tRNA pools can reflect adaptation to demanding protein synthesis needs and/or codon usage, the variations in the expression of the individual tRNA isodecoders are documented but poorly understood. Such expression variations are hypothesized to contribute to non-canonical tRNA functions, yet physiological relevance remains ambiguous. We report here that specific tRNA isodecoders can be functionally repurposed through cleavage that produces tRNA-derived RNAs (tDRs). The repurposing employs nucleotide variations in isodecoders leading to the production of distinct sets of tDRs with variable bioactivities.

## Introduction

tRNA is an adaptor RNA molecule fundamental to protein synthesis. tRNAs are the most abundant and diverse class of RNA species present in eukaryotes. The human genome encodes approximately 500 putative tRNA genes (1). The number of tRNA genes significantly exceed the number of distinct tRNAs required for mRNA translation (30-50 depending on species) by >10 fold. Such genetic excess is evolutionarily conserved from bacteria to humans, and high copy number of tRNA genes is required for abundant tRNA production during periods of enhanced proliferation and growth. Biogenesis of tRNA occurs in the nucleus, where it is transcribed by RNA Pol III and undergoes several rounds of enzymatic cleavage to form mature tRNA from its precursor form. Once in the cytoplasm, a tRNA molecule undergoes aminoacylation that involves enzymatic addition of amino acid at its 3’-end. Interestingly, two tRNA molecules with different sequences at the anti-codon loop may accept the same amino acid and are known as isoacceptors. In turn, isodecoders are different tRNA molecules with same anticodon sequence with minor sequence differences elsewhere in the body (2).

Published work suggests that all these minor changes in isodecoders are well tolerated for interaction with the ribosome and for aminoacylation since sequence changes do not cause any distortions in tRNA canonical secondary and tertiary structures (3) (4) (5). Overall, while the presence of tRNA isodecoders adds complexity to the understanding of translation regulation, further research is needed to fully elucidate their biological significance and mechanisms of action.

The presence of isoacceptor tRNAs is intrinsically connected to mRNA codon bias, and hence adds an extra layer of heterogeneity in regulating protein synthesis in vertebrates. In agreement with this, there are reports of tissue-specific expressions of distinct tRNA genes in vertebrate organisms (6). However, the biological function of the presence of tRNA isodecoders towards regulating canonical and non-canonical function of tRNA remains ambiguous. Earlier work from the Pan lab, based on the analysis of the efficiency of tRNA isodecoders in stop-codon suppression in human cell lines, suggested that specific isodecoder tRNAs display a large difference in their activities (7). These suppression activities within sets of isodecoders were ranging up to -20 fold despite similar stabilities and levels of aminoacylation *in vivo*. This work has also suggested that sequence changes within the tRNA molecule itself may enhance its non-canonical function by potentially reducing competition from the translation apparatus (7). Recently, number of non-canonical functions for tRNAs are reported. They range from tRNA functions in retroelement biology, stress adaptation, transcription and splicing, regulation of protein stability as well as immune responses (8).

Interestingly, the genetic excess of tRNA genes is also intrinsic to another evolutionary conserved process, tRNA cleavage. tRNA variants exist to serve as precursors to novel tRNA species (tRNA-derived RNAs, tDRs) (9) that mediate processes, beyond their canonical role in protein synthesis. These tDRs are produced through regulated tRNA cleavage by specific ribonucleases (RNases). We previously discovered that challenging human cells with a variety of abiotic stresses activates RNase angiogenin (ANG), which cleaves anticodon loops of tRNAs to produce specific class of tDRs, called tRNA-derived stress-induced <u>RNAs</u> (tiRNAs) (10). Growing evidence suggests that several tDRs engage in mRNA translation regulation, with a subset of tiRNAs acting as translation silencers that specifically inhibit mRNA translation initiation. In contrast to other small non-coding RNAs (ncRNAs) that recognize their targets via base pairing, tiRNA-mediated repression does not rely on sequence complementarity, and thus constitutes a novel mechanism of ncRNA-mediated target mRNA recognition/regulation. Specifically, terminal oligopyrimidine (TOG) motif containing tiRNAs such as 5’-tiRNA^Ala-AGC^ and 5’-tiRNA^Cys-CGA^ form G-quadruplex structures (G4-tiRNAs) (11) (12). These G4-tiRNAs inhibit canonical (cap-dependent) translation initiation but do not target non-canonical Internal Ribosome Entry Site (IRES)-driven translation initiation. The same subset of 5’-tiRNAs (derived from tRNAs^Ala/Cys^) also promotes the assembly of Stress Granules (SGs) (13), RNA granules with pro-survival and stress-adaptive functions, which help cells adapt to adverse environmental conditions and thus enhance their survival.

In addition to G4-tiRNAs, other tDRs are also implicated in translational control [reviewed in (14) (15)]. 5’tsGln is reported to regulate ribosomal protein translation by interacting with the multi-synthetase complex (16). 5’-tiRNA^Pro^ can associate with ribosomes and function as translation repressor in mammalian cells (17). On the other hand, in human parasite *T. brucei*, 3’tDR^Thr^ can stimulate global translation upon stress recovery by associating with the ribosomes (17). While reports of several tRNA derived fragments in translation regulation are growing with time, the molecular mechanism of their action remains poorly understood.

Expanding on our previous data, we hypothesized that different isodecoder tRNA fragments produced upon cleavage or maturation, may have different non-canonical functions. We propose that sequence variability observed in isodecoder tRNAs could be a means of fine-tuning the regulatory roles of tRNA fragments in various molecular processes. In our study, we screened and found specific isodecoders of 5’-tiRNA^Gly-GCC^ and 5’-tiRNA^His-GTG^ to possess translation repression activity. In addition, tRNA^His-GTG^ can be cleaved by ANG and RNase L (18) to produce different fragment length tiRNAs, each possessing translation repression activity. We show that single-nucleotide variation at the 5’-end of 5’-tiRNA^Gly-GCC^ and 5’-tiRNA^His^-^GTG^ dictates efficiency of translation repression activity for reporter mRNAs. Similarly, we tested translation repression activity of different 5’-tiRNAs derived from tRNA^Cys-GCA^ isodecoders that differ in the number of guanosine residue present in the TOG motif. Our result suggests that single nucleotide variation in tiRNAs derived from different isodecoders can modulate the translation repression activity of various tiRNAs.

## Results

### Non-G4-assembling 5’-tiRNAs repress mRNA translation

Previous reports from our lab demonstrated functions of G4-tiRNAs in repressing mRNA translation initiation. Mechanistically, this is achieved by targeting the HEAT domain of translation initiation factor eIF4G within cap-binding eIF4F complex (19). Here, we further expand our analysis towards other 5’-tiRNAs. Specifically, we tested the effect of the presence of different synthetic 5’-tiRNAs on translation of *Renilla luciferase* reporter mRNA using *in vitro* translation systems (IVTs) based on different mammalian extracts (**Fig. 1A, Supp. Fig. S1B**).

**Figure 1.**
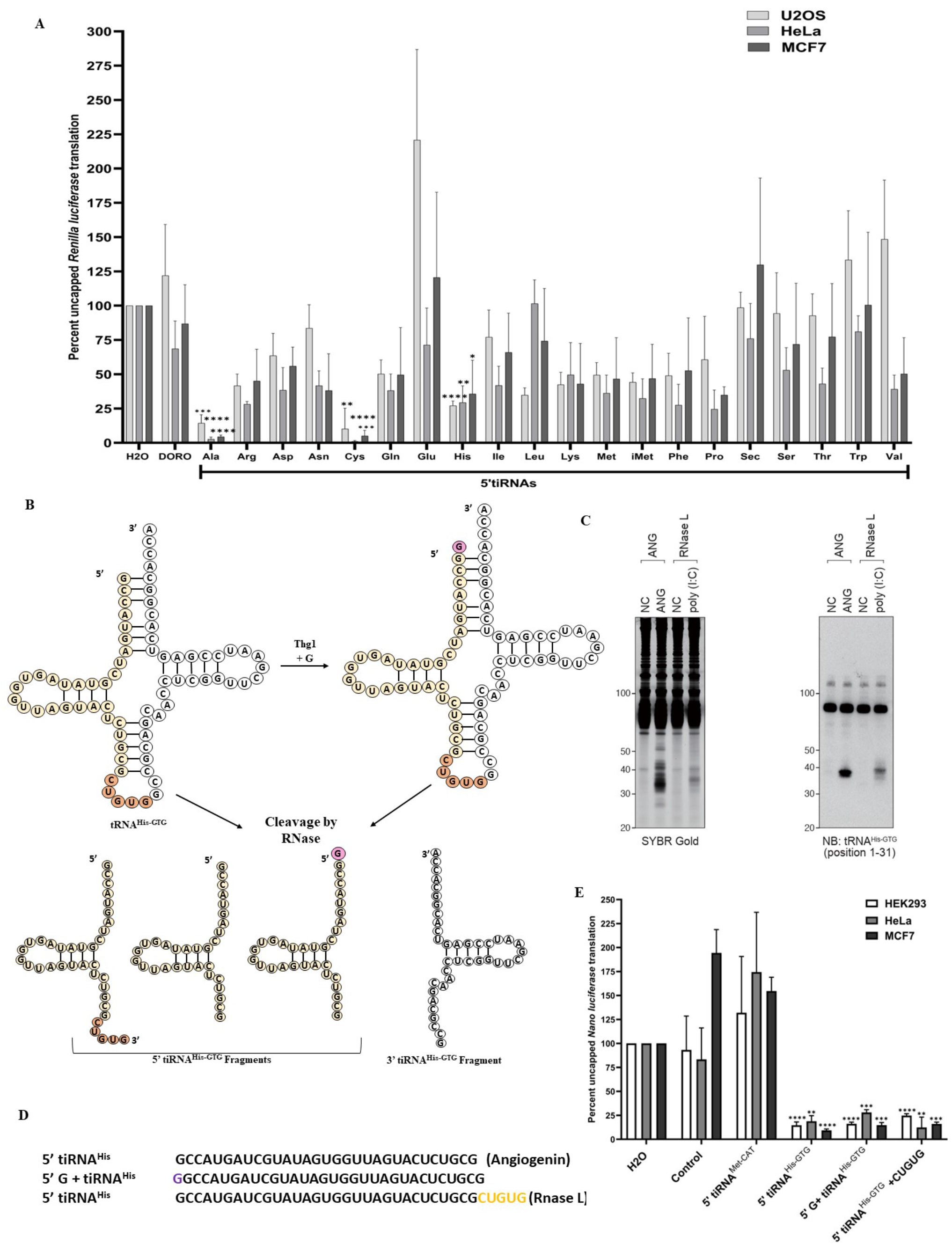
Screening 5’-tiRNAs for translation repression function. A: *In vitro* translation of uncapped *Renilla* luciferase reporter mRNA based on different mammalian cell extracts in the presence of various synthetic 5’-tiRNAs. B: Schematics on of tRNA^His-GTG^ -derived fragments produced by different cleavage. C: SYBR gold-stained gel (left) of total RNA isolated from U2OS cells and northern blot (right) probed against tRNA^His-GTG^ shows fragments generated after Angiogenin treatment or transfection with poly (I:C) (RNase L induction). D: Sequences of different 5’-tiRNA^His^ taken for *in vitro* translation assays in (E). E: *In vitro* translation assay of different 5’-tiRNA^His-GTG^ fragments in HEK293 translation extracts using capped *Nano luciferase* reporter mRNA.

As reported earlier, we observed 5’-tiRNA^Ala-AGC^ and 5’-tiRNA^Cys-GCA^ inhibiting mRNA translation in all three mammalian extracts used (**Fig. 1A**). Interestingly, we also observed that 5’-tiRNA^His-GTG^ represses translation of reporter mRNAs in different mammalian extracts (**Fig. 1A**, of uncapped *Renilla luciferase;* **Fig 1E**, *uncapped Nano luciferase; Fig. S1A, capped Nano luciferase)*. Unlike G4-tiRNAs, 5’-tiRNA^His-GTG^ does not harbor TOG sequence (see multi-sequence alignment of various tiRNAs used in the study, *Supp. Figure S1A*). Hence, we hypothesized that 5’-tiRNA^His-GTG^ may repress mRNA translation in a manner different from 5’-tiRNA^Ala-AGC^.

### Different 5’tiRNA^His-GTG^ variants possess translation repression activity

tRNA^His^ is unique in that it is a target of both RNase L and ANG, thereby producing 5’ tiRNA fragments of different length (**Fig. 1B-C**). In the same time, 5’-tiRNA^His^ significantly inhibits translation of mRNA reporters (**Fig. 1A, Fig.1E and Fig S1A**). Combining these two observations, we reasoned that tRNA^His^ would be a suitable model to test if slight variations in tRNA sequence (as it is seen with isoacceptors and isodecoders) or length can dictate variations in their non-canonical functions, upon tRNA cleavage. A unique feature of tRNA^His-GTG^ is the presence of an extra G residue at -1 position (_-1_G) on its 5’-end, which is added post-transcriptionally in eukaryotes by the enzyme THG1 (tRNA^His^ guanylyl transferase) (**Fig. 1B**). Consequently, it is reported that variants of 5’-tiRNA^His-GTG^ exist with or without (_-1_G) on its 5’-end (20). In addition, 5’-tiRNAs generated after cleavage of tRNA^His-GTG^ have different length depending upon the cleavage enzyme (**Fig. 1B-D**). Specifically, ANG-mediated cleavage results in a shorter variant of 5’-tiRNA^His-GTG^ than variant produced by RNase L cleavage (21). We tested whether different length variants of 5’-tiRNA^His-GTG^ affect their translation repression functions using IVTs based on HEK293, HeLa and MCF7 extracts (**Fig. 1E**). Our data based on IVT systems with capped *Nano luciferase* reporter mRNA revealed that the different fragment length 5’-tiRNA^His-GTG^ could inhibit translation to an equal extent (**Fig. 1E**). Based on these results, we decided to do further downstream assays with 5’-tiRNA^His-GTG^ lacking the extra G at - 1 position since they are reported to be more abundant in cells (20).

### Single nucleotide variation in 5’-tiRNA^His-GTG^ derived from a specific isodecoder imparts translation repression function

In humans, tRNA^His^ has only one isoacceptor, tRNA^His-GTG^, and is encoded by ten isodecoder genes. tRNA^His-GTG-2-1^ (further referred as 2-1) differs from the other nine genes by the presence of adenine at the position 4 of the amino acid acceptor arm while all other tRNA isodecoders (1-1 to 1-9) contain guanosine (**Fig. 2A**). Consequently, post-RNase cleavage of 2-1 isodecoder tRNA, this adenosine is present in the corresponding the 5’-tiRNA^His-GTG^ variant.

**Figure 2.**
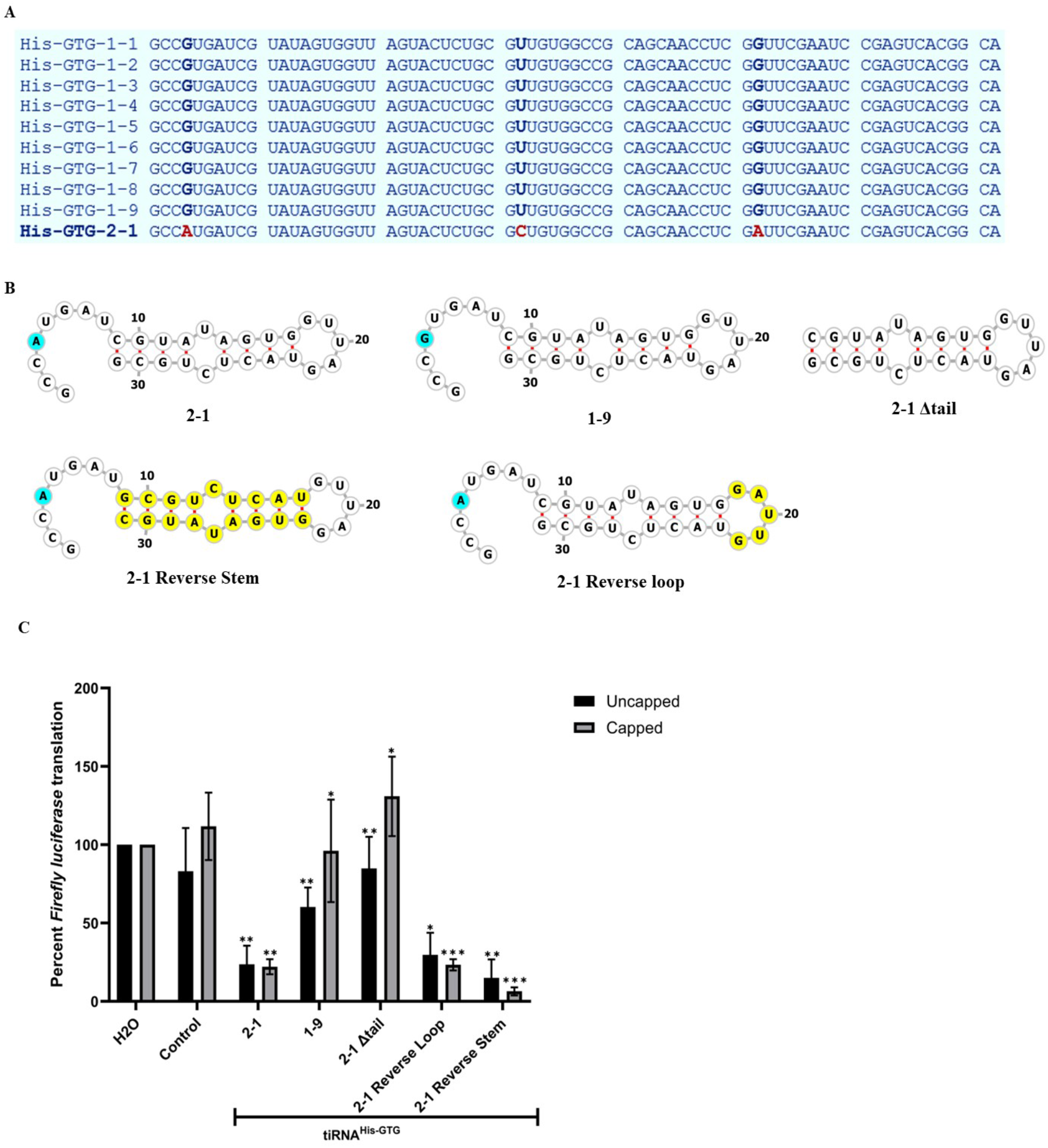
Isodecoder-specific variants of tiRNA^His-GTG^ possess translation repression activity. A: Multi-sequence alignment of various isodecoders of tRNA^His-GTG^ using Mult-Alin software tool. Amongst the 10 isodecoders present in humans, only tRNA^His-GTG^ 2-1 has an “AUG” with adenine at the position 4 of the 5’-end. B: Secondary structure prediction of 5’-tiRNA^His-GTG^1-9, 2-1 and mutants examined in *in vitro* translation assay. C: *In vitro* translation assay of capped and uncapped *Firefly* luciferase reporter mRNAs in HEK293-based translation extracts in the presence of wild-type and different mutants of 5’-tiRNA^His-GTG^ (p-value are shown for 2-1 as compared to water control, 1-9 and 2-1Δtail is with respect to 2-1, and 2-1 reverse-loop and reverse-stem is with respect to water control, n=3).

We decided to test if the ability of 5’-tiRNA^His-GTG^ to repress translation is contingent on the presence of G or A in the sequence. We used synthetic forms of 5’-tiRNA^His-GTG^ isodecoder variants containing G or A at the position 4 of the tiRNA (**Fig. 2B**). Secondary structure prediction using the software RNAfold suggests that A-G substitution does not affect a putative tiRNA folding. Surprisingly, we observed that A to G substitution within 5’-tiRNA^His-GTG^ abolished its ability to repress translation of both capped and uncapped *Firefly luciferase* mRNAs (**Fig. 2C**). In addition, our structure-functional analysis of mutant 5’-tiRNA 2-1 variants, which lack 5’-overhanging tail (2-1 Δtail) or flipped stem or loop structures (2-1 Reverse Stem and 2-1 Reverse Loop, correspondingly) suggest that the presence of A in the 5’-tail sequence is indispensable for translation repression function for both capped and uncapped reporters (**Fig. 2 B-C**).

### Single nucleotide variations in other 5’-tiRNAs can affect their structure and function

Previous report from our lab has shown that 5’-tiRNA^Gly-GCC^ (both synthetic and isolated from cells) represses reporter mRNA translation in rabbit reticulocyte lysate (22) (23). Interestingly, a common feature of tRNA^Gly-GCC^ and tRNA^His-GTG^ is the presence of single nucleotide variation within an “AUG” sequence at the 5’-end, predicted to be single-stranded (**Fig. 3A-B**). This observation encouraged us to further examine different isodecoders of tRNA^Gly-GCC^ via multi-sequence alignment and analyze translation repression function between 5’-tiRNA variants containing single nucleotide variations (**Fig. 3**., and **Supp. Fig. S1A**).

**Figure 3.**
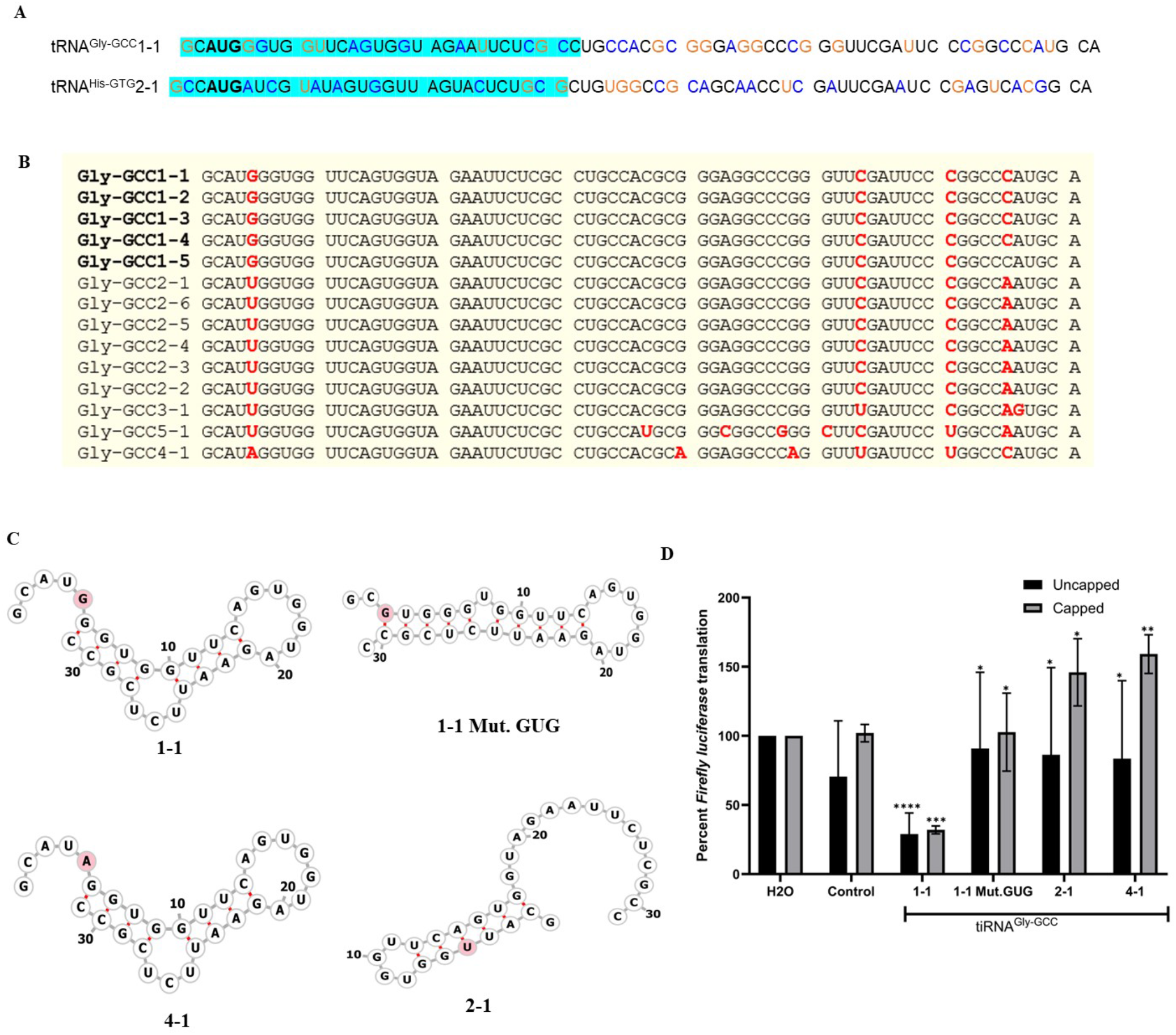
Structure-functional similarities between 5’-tiRNA^Gly-GCC^ 1-1 and 5’-tiRNA^His-GTG^ 2-1 responsible for translation repression activities. A: Sequence alignment of tRNA^Gly-GCC^ and tRNA^His-GTG^ The region highlighted by blue color is the product generated post angiogenin cleavage (5’-tiRNA). B: Multi-sequence alignment of various tRNA^Gly-GCC^ isodecoders using Mult-Alin online software tool. Amongst the 14 isodecoders present in humans, five isodecoders of tRNA^Gly-GCC^ 1-1 to 1-5 harbor an “AUG” sequence at the 5’-end. C: Secondary structure prediction of different isodecoders of 5’-tiRNA^Gly-GCC^ using RNA-Fold software. D: *In vitro* translation assay of capped and uncapped *Firefly luciferase* reporter mRNA in HEK293 translation extracts in the presence of different isodecoder-specific variants of 5’-tiRNA^Gly-GCC^.

In humans, there are three isoacceptors for tRNA^Gly^, namely tRNA^Gly-GCC^, tRNA^Gly-CCC^ and tRNA^Gly-TCC^. The tRNA^Gly-GCC^ further has 14 isodecoders (**Fig. 3B**) amongst which tRNA^Gly-GCC^ 1-1, 1-2, 1-3, 1-4 and 1-5 have an “AUG” hallmark sequence at the 5’-end. This 5’-end AUG sequence is similar to that observed in 5’-tiRNA^His-GTG^, a feature that differentiates the five isodecoders of tRNA^Gly-GCC^ from the other nine. We used synthetic forms of 5’-tiRNA^Gly-GCC^ 1-1, 4-1 and 2-1 (which harbor G, U and A nucleotide variations at the position 5 of their 5’-end sequences) and tested their ability to repress translation of mRNA reporters in HEK293-derived IVT. We observed that 5’-tiRNA^Gly-GCC^ 2-1 and 4-1 could not repress translation of either capped or uncapped *Firefly* luciferase reporter mRNA translation owing to the presence of U and A nucleotides instead of G at the 5’-end of the 1-1 variant (**Fig. 3D**). Interestingly, unlike tRNA^His-GTG^, single nucleotide variation in the 2-1 isodecoder tRNA^Gly-GCC^ is also predicted to affect its secondary structure where 5’-end is base-paired while compared to the 5’ single-stranded structure of variants 1-1 and 4-1 (**Fig. 3C**). Further mutational analysis that converts a putative “AUG” motif to “GUG” (substitution of A to G at the position 3, 1-1 Mut.GUG variant (**Fig. 3C**)) also affects predicted secondary structure of 1-1 variant and, consequently, inactivates its repressor activity towards mRNA reporters.

We further extended our analysis on the significance of single nucleotide variation affecting translation repression activities to TOG motif containing tiRNAs. Multi-sequence alignment of different isodecoders of tRNA^Cys-GCA^ suggested the presence of variations in the number of guanosine residues in the TOG motif in tRNA^Cys-GCA^ 17-1 and 20-1 (**Supp. Fig. S2A**). Secondary structure predictions indicated that 2-1 and 17-1 variants of tRNA^Cys-GCA^ form similar structures with 5’-single-stranded overhang while 20-1 variant would fold in a different structure (**Supp. Fig. S2B**). IVT assay using HEK293 extract revealed that the presence of 5 Gs versus 4Gs in the TOG motif of 5’-tiRNA^Cys-GCA^ (2-1 versus 17-1 variants) moderately affects its ability to repress translation of uncapped mRNA reporter but loses its ability to repress capped mRNA (**Supp. Fig. S2C**). At the same time, 5’-tiRNA^Cys-GCA^ 20-1 variant losses its ability to repress translation of both mRNA reporters (**Supp. Fig. S2C**). Surprisingly, we observed that ability to repress reporter mRNA does not coincide with the ability of 5’-tiRNA^Cys-GCA^ variants to form G-quadruplex structure as observed by NMM-in-gel staining (**Supp. Fig. S2D**). In fact, 5’-tiRNA^Cys-GCA^ 20-1 variant can assemble G4 structures although its 5’-end contains C to G substitution at the position 4 of the TOG motif.

## Discussion

The functional relevance of diversity in tRNA genes since its discovery has long been a paradox in the field (24). In single cellular organisms such as bacteria and yeast, the occurrence of isodecoder tRNA genes is less common. For example, there are reportedly 51 isodecoder genes out of a total of 274 tRNA genes in budding yeast. In contrast, in humans, the number of isodecoder genes is as high as 274 out of a total of 446 tRNA genes (25). In humans, cell-specific transcription of distinct set of tRNA isodecoder gene helps in gaining tissue level complexity (25), (6), (26). Importantly, the work from the Ackerman lab showed that the regulation of individual isodecoder tRNAs enables translational control, and a pathogenic mutation in one of 5 isodecoders of the tRNA^Arg-UCU^ underlines neurodegenerative phenotype in central nervous system (27).

From the perspective of canonical function of tRNA, presence of specific isodecoder and isoacceptor sets can correlate with codon bias and hence modulate the translation efficiency of different mRNAs (**Fig. 4A**, canonical function). From the perspective of non-canonical functions (**Fig. 4A**, non-canonical function), expression of different isodecoder tRNAs can further diversify the cellular population of tDRs and hence functions associated with them. The molecular impact of the presence of distinct tDRs towards cellular functions is increasingly appreciated yet remains poorly understood. Recent data also suggest that changes in the abundance of mature tRNAs are not directly reflected by differential tRNA gene expression unlike changes in tDR levels (28).

**Figure 4.**
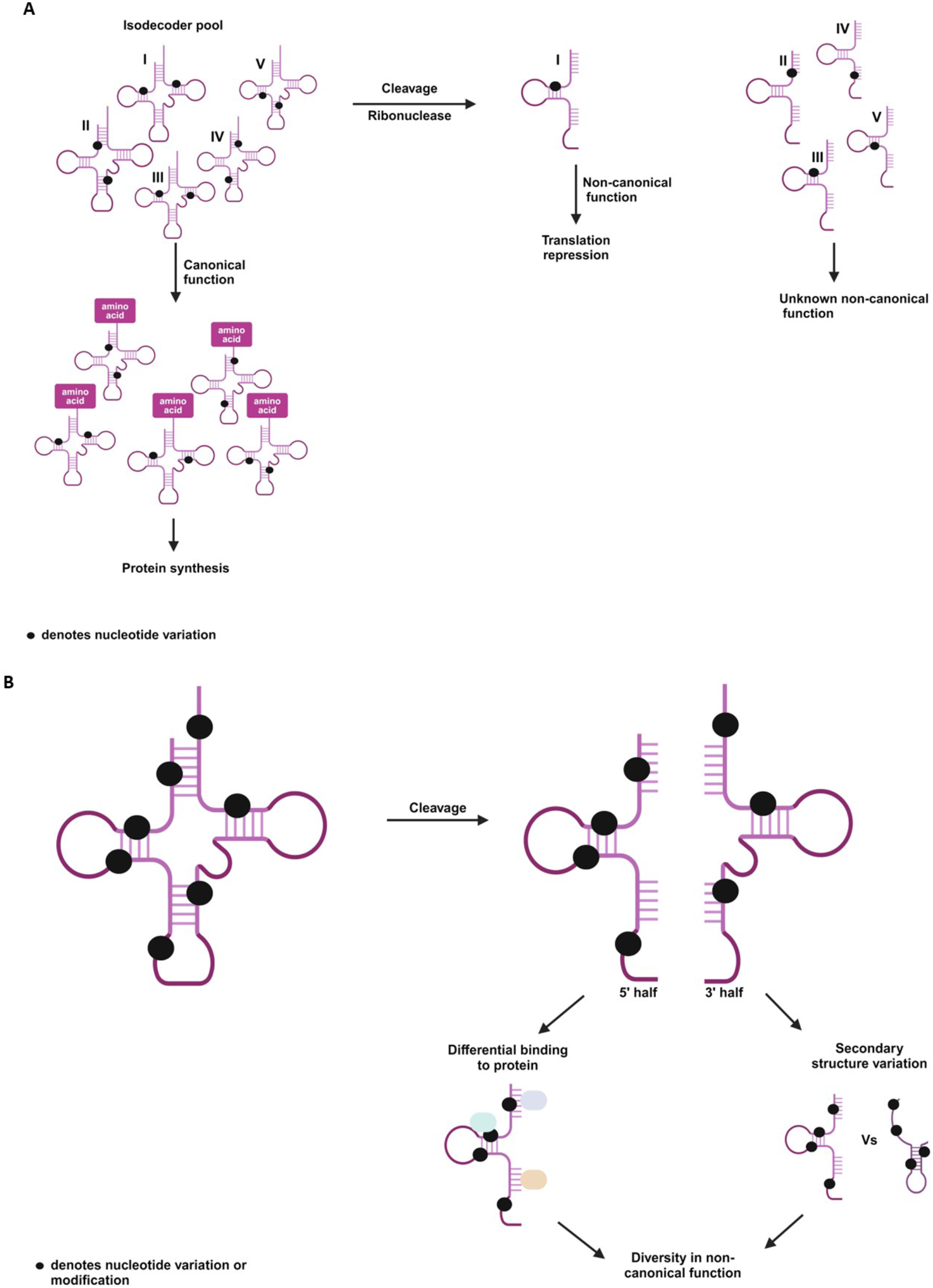
Proposed model. A: tRNA isodecoders are molecules that have same anti-codon sequence and hence are charged by the same amino acid but harbor sequence variation elsewhere in their bodies, as depicted by the black dot(s). Upon cleavage, sequence variation within isodecoders’ bodies can affect its ability to perform non-canonical function such as translation repression as described in our study. B: Presence of isodecoder tRNA with specific base modification can diversify the pool of tRNA derived fragments in a cell. Upon cleavage, a given isodecoder tRNA fragment could attract specific set of proteins or fold differently, thereby participating in several non-canonical functions providing an additional level of complexity reflected by some biological functions.

Here, we provide evidence that diversity of tRNA isodecoders’ functions can be further expanded via tRNA cleavage. RNase-mediated cleavage of different tRNA isodecoders can generate tDRs that vary in their length, nucleotide composition or modification status. Our model suggests that in turn, such variable tDRs derived from the subsets of isodecoder tRNAs (with or without specific base modification) can assemble into different structures and conduct different functions (**Fig. 4B**). Our *in vitro* studies suggest that even minimal single nucleotide change within tiRNAs can dramatically affect its non-canonical functions, e.g. in the ability to repress mRNA translation.

5’-tiRNA^Gly-GCC^ (1-5 variant) is a highly abundant tDR found to be expressed in hippocampus hematopoietic and lymphoid tissues but depleted in gonadal tissues (29), (30), (31). Moreover, it is known to be also present in extracellular serum where it circulates as a stable complex as opposed to enclosed vesicle (30). The functional relevance of the tissue specific expression of 5’-tiRNA^Gly-GCC^1-5 and its mechanism of action remains elusive. It will be important to test how translation repression function of 5’-tiRNA^Gly-GCC^1-5 helps in maintaining cellular homeostasis in hippocampus, hematopoietic and lymphoid tissues.

The endogenous 5’-tiRNA^Gly-GCC^ is more active as a translation inhibitor than its synthetic form, emphasizing the potential role of its nucleotide modification towards regulating mRNA translation (23). Interestingly, the 5’-tiRNA^Gly-GCC^ 2-1 variant has been reported to form homodimer or heterodimer with 5’-tiRNA^Gln^ (19) (32). Secondary structure prediction using RNAfold software suggests that 5’-tiRNA^Gly-GCC^2-1 has significantly different folding pattern as compared to 5’-tiRNA^Gly-GCC^1-5 and 5’-tiRNA^Gly-GCC^ 4-1 (**Fig. 3C**). Unlike the 5’-tiRNA^Gly-GCC^ 2-1 variant, the “AUG” and “AUA” trinucleotides in 5’-tiRNA^Gly-GCC^1-5 and 5’-tiRNA^Gly-GCC^4-1 are predicted to not base pair. We speculate that it may attract unique and unidentified protein binding partners to carry out non-canonical functions (as in the model, **Fig 4B**). It will be important to probe whether different secondary structure pattern of 5’-tiRNAs derived from distinct isodecoders of tRNA^Gly-GCC^ could affect their binding partners and hence non-canonical function. It should be noted that while 5’-tiRNA^Gly-GCC^ 2-1 and 4-1 fragments do not modulate translation of reporter mRNAs tested in our study, we speculate that they may have other, mRNA translation unrelated, non-canonical functions yet to be identified.

Our data show that other isodecoders of 5’-tiRNA^His-GTG^ that harbor G instead of A nucleotide at the position 4 of 5’-end cannot repress translation (**Fig. 2B-C**). The G-containing 5’-tiRNA^His-GTG^ has been implicated in the progression of colon cancer (33). Mechanistically, it promotes the expression of anti-apoptotic and pro-proliferative genes by targeting large tumor suppressor kinase2, thereby inactivating the Hippo signaling pathway. Similarly, *Mycobacterium bovis* infection has been reported to elevate the levels of 5’-tiRNA^His-GTG^ derived from G-encoding isodecoder in human monocyte-derived macrophages. During infection, these tiRNAs are packaged in extracellular vesicles and delivered into the recipient cells where they function to activate the innate immune cell response via stimulation of TLR7 (34). On the other hand, 5’-tiRNA^His-GTG^ 2-1 is less characterized tDR, with expression of parental tRNA^His-GTG^ 2-1 limited specifically to pancreas and neural tissues (personal communication of Prof. Todd Lowe, UCSC). Hence, the functional and disease relevance of the presence of 5’-tiRNA^His-GTG^ 2-1 variant in specific tissue needs further investigation.

In summary, our study shows that tRNA cleavage can repurpose a single nucleotide difference in different isodecoders of tRNA^His-GTG^, tRNA^Gly-GCC^ and tRNA^Cys-GCA^ towards non-canonical function. We propose that tRNA diversity underlines multiple hidden layers of gene expression regulation operated via non-canonical tRNA functions.

## Materials and methods

### Tissue culture and treatment

Wild-type U2OS, HEK293, Hela and MCF7 cells were grown at 37°C moist incubator with 5% CO_2_ in Dulbecco’s modified Eagle’s medium (DMEM) supplemented with 10% fetal bovine serum (FBS) (Sigma) and 1% of penicillin/streptomycin (Sigma).

For tDR production, U2OS cells were treated with recombinant human Angiogenin (R&D systems) at 0.5 µg/mL for 1 h or transfected with 2 µg/mL Poly (I:C) HMW (Invivogen) for 6 h using Lipofectamine 3000 reagent (Thermo Fisher) according to the manufacturer’s instructions.

### *In vitro* transcription and capping

The T7-Nluc-control and T7-pHRL-control plasmids were isolated and subjected to overnight digestion by Sac II (NEB, R01575). The digested plasmids were extracted using Phenol:Chloroform:IsoamylAlcohol (25:24:1) solution. Typically, 200-500ng of digested plasmid was used for transcription using Promega Ribomax T7 transcription kit (cat ID#P1320) as per manufacturer’s protocol. Post transcription, the RNA was purified by first passing it through G25-sephadex column, Amersham (to remove unincorporated nucleotides) and then the eluent was further purified using NEB-monarch RNA cleanup kit (cat ID #T2040L). The quality of the transcribed reporter mRNA was checked in 6% TBU gel and then stored at -80 to be used for *in vitro* translation reactions.

*Firefly luciferase* reporter mRNA was purchased from Promega and capped using cell-script capping kit (cat ID# CSCCS1710) as per manufacturer’s protocol.

### Preparation of translation competent mammalian extract

After reaching desirable confluency, mammalian cells were detached via trypsinization at 37°C for 2min. Equal volume of DMEM medium was added to stop the trypsinization reaction and cells were pelleted at 1000rpm for 2min. The media was removed, and cells were suspended in 1ml of hypotonic lysis buffer (10mM HEPES pH7, 10mM potassium acetate, 0.5mM magnesium acetate, 5mM DTT and one tablet of mini EDTA free protease inhibitor tab from Roche). The cells were transferred to a 1.5ml microcentrifuge tube and centrifuged at 1000 RCF at 4°C for 1min. The supernatant was removed, and the cells were resuspended in chilled lysis buffer (volume of lysis buffer is same as volume of packed cells). Cells were tumbled at 4°C for 1hr followed by homogenization by passing through a 1ml 27G needle syringe for 10-12 times. The resulting lysate was centrifuged at 14000 RCF for 1 min and 40ul of supernatant was aliquoted in 1.5ml microcentrifuge tubes. The tubes were snap frozen in dry ice followed by immediate storage at -80°C.

### NMM staining

Detection of G-quadruplex structure was done as described in (35). Briefly, synthetic 5’tiRNA isodecoders of tiRNA^Cys^ were folded in TE buffer with 150mM KCl. Post folding the samples were run in 20% native TBE gel and stained with 1µM NMM followed by observation under UV light. The same gel was destained for 30min in TBE buffer and then imaged after 2min of EtBr staining.

### Northern blotting

Northern blotting was performed as we previously reported (36). The probe for 5’-tiRNA^His-GTG^ (5’-CGCAGAGTACTAACCACTATACGATCACGGC-3’) was synthesized by IDT.

### *In vitro* translation

*In vitro* translation reaction in mammalian extracts were performed as described in previous report (37). Briefly, a 10µl reaction was set up with 10pm of appropriate tiRNAs, 100-150ng of reporter mRNA, 1.25mM of magnesium acetate, 150mM potassium acetate, 1X translation buffer (having 1.6mM HEPES pH7, 2mM creatine phosphate, 0.01 µg/ml creatine kinase, 10 µM spermidine and 10 µM L-amino acid mix, cat ID # L4461), 1 unit of RNase inhibitor (Promega, cat ID# N251B) and micrococcal nuclease (Thermo Fisher, cat ID# 88216) treated mammalian extract. The tubes were then incubated at 37°C for 1hr followed by taking luminescence reading. Depending upon the reporter mRNA used, 20-50% of the reaction was mixed with either Firefly, Renilla or Nano-luciferase reagent and luminescence was measured using Promega Glomax Explorer Machine.

## Acknowledgments

We deeply appreciate Yuichiro Adachi, Claire Riggs, and Prakash Kharel for their technical expertise, comprehensive knowledge, and protocol support, along with their assistance in data analysis and Victoria Ivanova, Allison Williams, and Safiyah Zubair for supplying experimental reagents, offering technical insights and support.

This work was supported by funds from National Institutes of Health grant R35 GM126901 (P.J.A.), National Institutes of Health grant R01 GM126150 and R01 GM146997 (P.I.).

## Author Contributions

PI and PJA. designed research; NB, YA, YT, SML and VMA performed research; all authors analyzed data and provided helpful discussion; NB, VMA, PJA, PI.. wrote the paper. PJA, SML, YA and PI provided funding.

## Supplementary Information

### Supplementary Figure Legends

**Supplementary Figure S1.**
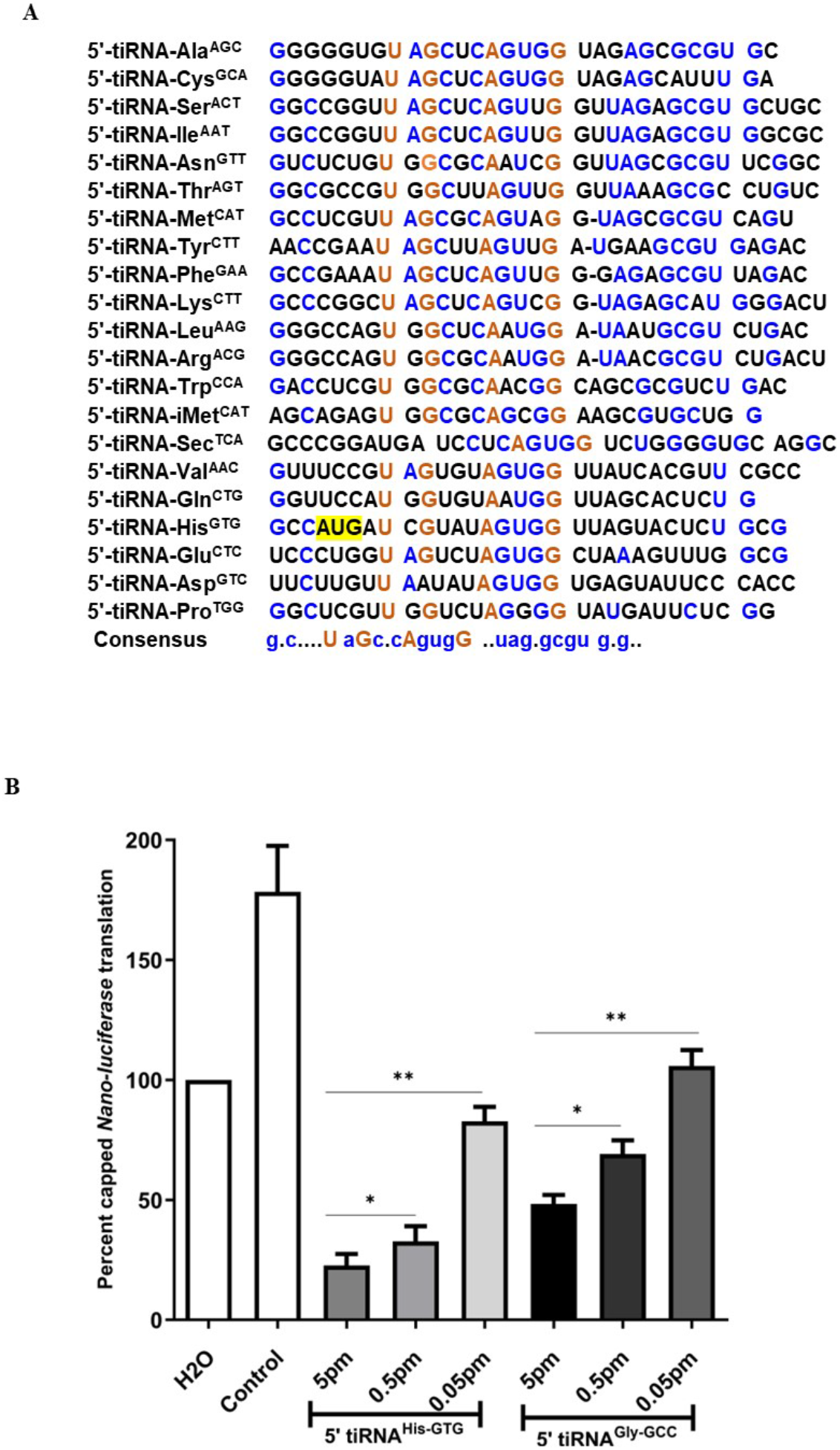
**(A)** Multi-sequence alignment of the 5’-tiRNAs used for *in vitro* translation assay. (**B)** Concentration-dependent translation repression of capped *Nano luciferase* reporter mRNA in HEK293-based translation extract by 5’-tiRNA^His-GTG^ and 5’-tiRNA^Gly-GCC^.

**Supplementary Figure S2.**
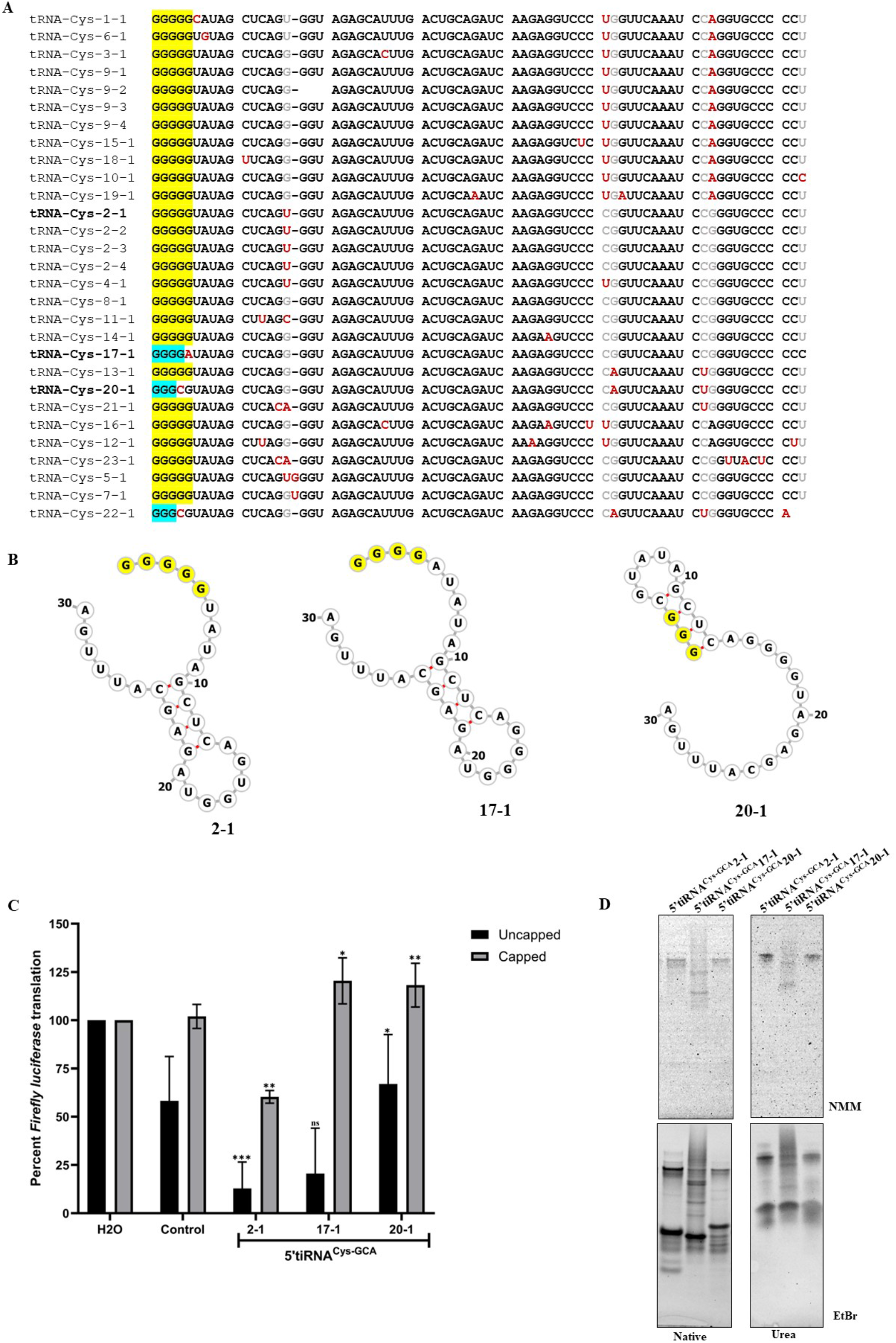
(**A)** Multi-sequence alignment of various isodecoders of tRNA^Cys-GCA^ using MultAlin software tool. Amongst the 29 isodecoders of tRNA^Cys-GCA^ present in humans, all isodecoders possess five guanosines at the 5’-end except tRNA^Cys-GCA^ (17-1, 20-1 and 22-1). (**B**) Secondary structure prediction (using RNAFold software) of different isodecoders of 5’-tiRNA^Cys-GCA^ used for *in vitro* translation assays. (**C**) *In vitro* translation assays of capped and uncapped *Firefly luciferase* reporter mRNAs in HEK293 translation extracts in the presence of different isodecoders of 5’-tiRNA^Cys-GCA^. (**D**) Different isodecoders of 5’-tiRNA^Cys^ run in a 20% TBE native gel or 15% urea gel after folding in KCl Buffer supporting G-quadruplex assembly. The gel was first stained with 1µm NMM (dye specific for parallel G4s) followed by destaining and staining with EtBr dye (general staining).

