## Supplementary Information for "Repurposing tRNA isodecoders for non-canonical functions via tRNA cleavage"

**Supplementary Figure S1**

**A**

| | |
|---|---|
| 5'-tiRNA-Ala$^{AGC}$ | GGGGGUGU AGCUCAGUGG UAGAGCGCGU GC |
| 5'-tiRNA-Cys$^{GCA}$ | GGGGGUAU AGCUCAGUGG UAGAGCAUUU GA |
| 5'-tiRNA-Ser$^{ACT}$ | GGCCGGUU AGCUCAGUUG GUUAGAGCGU GCUGC |
| 5'-tiRNA-Ile$^{AAT}$ | GGCCGGUU AGCUCAGUUG GUUAGAGCGU GGCGC |
| 5'-tiRNA-Asn$^{GTT}$ | GUCUCUGU GGCGCAAUCG GUUAGCGCGU UCGGC |
| 5'-tiRNA-Thr$^{AGT}$ | GGCGCCGU GGCUUAGUUG GUUAAAGCGC CUGUC |
| 5'-tiRNA-Met$^{CAT}$ | GCCUCGUU AGCGCAGUAG G-UAGCGCGU CAGU |
| 5'-tiRNA-Tyr$^{CTT}$ | AACCGAAU AGCUUAGUUG A-UGAAGCGU GAGAC |
| 5'-tiRNA-Phe$^{GAA}$ | GCCGAAAU AGCUCAGUUG G-GAGAGCGU UAGAC |
| 5'-tiRNA-Lys$^{CTT}$ | GCCCGGCU AGCUCAGUCG G-UAGAGCAU GGGACU |
| 5'-tiRNA-Leu$^{AAG}$ | GGGCCAGU GGCUCAAUGG A-UAAUGCGU CUGAC |
| 5'-tiRNA-Arg$^{ACG}$ | GGGCCAGU GGCGCAAUGG A-UAACGCGU CUGACU |
| 5'-tiRNA-Trp$^{CCA}$ | GACCUCGU GGCGCAACGG CAGCGCGUCU GAC |
| 5'-tiRNA-iMet$^{CAT}$ | AGCAGAGU GGCGCAGCGG AAGCGUGCUG G |
| 5'-tiRNA-Sec$^{TCA}$ | GCCCGGAUGA UCCUCAGUGG UCUGGGGUGC AGGC |
| 5'-tiRNA-Val$^{AAC}$ | GUUUCCGU AGUGUAGUGG UUAUCACGUU CGCC |
| 5'-tiRNA-Gln$^{CTG}$ | GGUUCCAU GGUGUAAUGG UUAGCACUCU G |
| 5'-tiRNA-His$^{GTG}$ | GCCAUGAU CGUAUAGUGG UUAGUACUCU GCG |
| 5'-tiRNA-Glu$^{CTC}$ | UCCCUGGU AGUCUAGUGG CUAAAGUUUG GCG |
| 5'-tiRNA-Asp$^{GTC}$ | UUCUUGUU AAUAUAGUGG UGAGUAUUCC CACC |
| 5'-tiRNA-Pro$^{TGG}$ | GGCUCGUU GGUCUAGGGG UAUGAUUCUC GG |
| Consensus | g.c....U aGc.cAgugG ..uag.gcgu g.g.. |

**B**

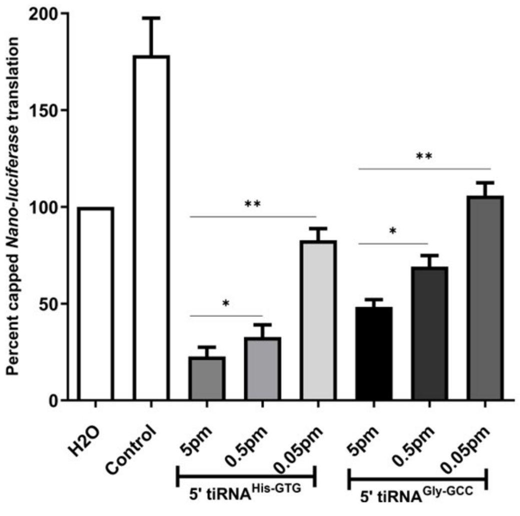

### Supplementary Figure S2

**A**

```
tRNA-Cys-1-1     GGGGGCAUAG CUCAGU-GGU AGAGCAUUUG ACUGCAGAUC AAGAGGUCCC UGGUUCAAAU CCAGGUGCCC CCU
tRNA-Cys-6-1     GGGGGUGUAG CUCAGU-GGU AGAGCAUUUG ACUGCAGAUC AAGAGGUCCC UGGUUCAAAU CCAGGUGCCC CCU
tRNA-Cys-3-1     GGGGGUAUAG CUCAGG-GGU AGAGCACUUG ACUGCAGAUC AAGAGGUCCC UGGUUCAAAU CCAGGUGCCC CCU
tRNA-Cys-9-1     GGGGGUAUAG CUCAGG-GGU AGAGCAUUUG ACUGCAGAUC AAGAGGUCCC UGGUUCAAAU CCAGGUGCCC CCU
tRNA-Cys-9-2     GGGGGUAUAG CUCAGG-    AGAGCAUUUG ACUGCAGAUC AAGAGGUCCC UGGUUCAAAU CCAGGUGCCC CCU
tRNA-Cys-9-3     GGGGGUAUAG CUCAGG-GGU AGAGCAUUUG ACUGCAGAUC AAGAGGUCCC UGGUUCAAAU CCAGGUGCCC CCU
tRNA-Cys-9-4     GGGGGUAUAG CUCAGG-GGU AGAGCAUUUG ACUGCAGAUC AAGAGGUCCC UGGUUCAAAU CCAGGUGCCC CCU
tRNA-Cys-15-1    GGGGGUAUAG CUCAGG-GGU AGAGCAUUUG ACUGCAGAUC AAGAGGUCUC UGGUUCAAAU CCAGGUGCCC CCU
tRNA-Cys-18-1    GGGGGUAUAG UUCAGG-GGU AGAGCAUUUG ACUGCAGAUC AAGAGGUCCC UGGUUCAAAU CCAGGUGCCC CCU
tRNA-Cys-10-1    GGGGGUAUAG CUCAGG-GGU AGAGCAUUUG ACUGCAGAUC AAGAGGUCCC UGGUUCAAAU CCAGGUGCCC CCC
tRNA-Cys-19-1    GGGGGUAUAG CUCAGG-GGU AGAGCAUUUG ACUGCAAAUC AAGAGGUCCC UGAUUCAAAU CCAGGUGCCC CCU
tRNA-Cys-2-1     GGGGGUAUAG CUCAGU-GGU AGAGCAUUUG ACUGCAGAUC AAGAGGUCCC CGGUUCAAAU CCGGGUGCCC CCU
tRNA-Cys-2-2     GGGGGUAUAG CUCAGU-GGU AGAGCAUUUG ACUGCAGAUC AAGAGGUCCC CGGUUCAAAU CCGGGUGCCC CCU
tRNA-Cys-2-3     GGGGGUAUAG CUCAGU-GGU AGAGCAUUUG ACUGCAGAUC AAGAGGUCCC CGGUUCAAAU CCGGGUGCCC CCU
tRNA-Cys-2-4     GGGGGUAUAG CUCAGU-GGU AGAGCAUUUG ACUGCAGAUC AAGAGGUCCC CGGUUCAAAU CCGGGUGCCC CCU
tRNA-Cys-4-1     GGGGGUAUAG CUCAGU-GGU AGAGCAUUUG ACUGCAGAUC AAGAGGUCCC UGGUUCAAAU CCGGGUGCCC CCU
tRNA-Cys-8-1     GGGGGUAUAG CUCAGG-GGU AGAGCAUUUG ACUGCAGAUC AAGAGGUCCC CGGUUCAAAU CCGGGUGCCC CCU
tRNA-Cys-11-1    GGGGGUAUAG CUUAGC-GGU AGAGCAUUUG ACUGCAGAUC AAGAGGUCCC CGGUUCAAAU CCGGGUGCCC CCU
tRNA-Cys-14-1    GGGGGUAUAG CUCAGG-GGU AGAGCAUUUG ACUGCAGAUC AAGAAGUCCC CGGUUCAAAU CCGGGUGCCC CCU
tRNA-Cys-17-1    GGGGAUAUAG CUCAGG-GGU AGAGCAUUUG ACUGCAGAUC AAGAGGUCCC CGGUUCAAAU CCGGGUGCCC CCC
tRNA-Cys-13-1    GGGGGUAUAG CUCAGG-GGU AGAGCAUUUG ACUGCAGAUC AAGAGGUCCC CAGUUCAAAU CUGGGUGCCC CCU
tRNA-Cys-20-1    GGGCGUAUAG CUCAGG-GGU AGAGCAUUUG ACUGCAGAUC AAGAGGUCCC CAGUUCAAAU CUGGGUGCCC CCU
tRNA-Cys-21-1    GGGGGUAUAG CUCACA-GGU AGAGCAUUUG ACUGCAGAUC AAGAGGUCCC CGGUUCAAAU CUGGGUGCCC CCU
tRNA-Cys-16-1    GGGGGUAUAG CUCAGG-GGU AGAGCACUUG ACUGCAGAUC AAGAAGUCCU UGGUUCAAAU CCAGGUGCCC CCU
tRNA-Cys-12-1    GGGGGUAUAG CUUAGG-GGU AGAGCAUUUG ACUGCAGAUC AAAAGGUCCC UGGUUCAAAU CCAGGUGCCC CUU
tRNA-Cys-23-1    GGGGGUAUAG CUCACA-GGU AGAGCAUUUG ACUGCAGAUC AAGAGGUCCC CGGUUCAAAU CCGGUUACUC CCU
tRNA-Cys-5-1     GGGGGUAUAG CUCAGUGGGU AGAGCAUUUG ACUGCAGAUC AAGAGGUCCC CGGUUCAAAU CCGGGUGCCC CCU
tRNA-Cys-7-1     GGGGGUAUAG CUCAGGUGGU AGAGCAUUUG ACUGCAGAUC AAGAGGUCCC CGGUUCAAAU CCGGGUGCCC CCU
tRNA-Cys-22-1    GGGCGUAUAG CUCAGG-GGU AGAGCAUUUG ACUGCAGAUC AAGAGGUCCC CAGUUCAAAU CUGGGUGCCC A
```

**B**

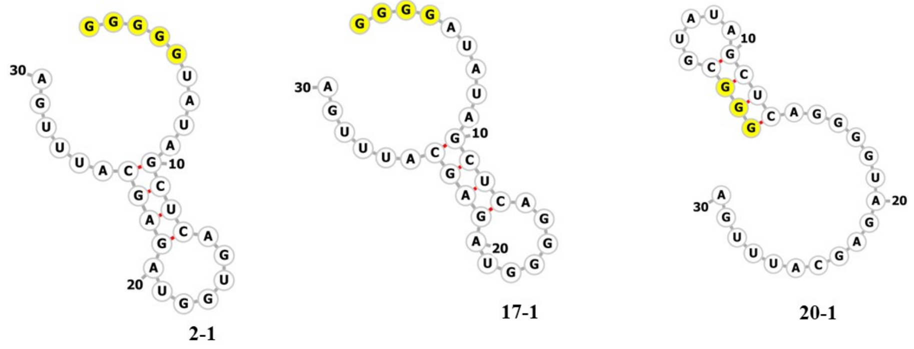


**C**

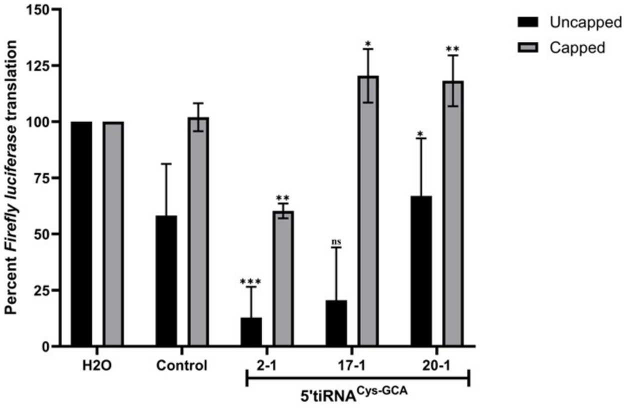


**D**

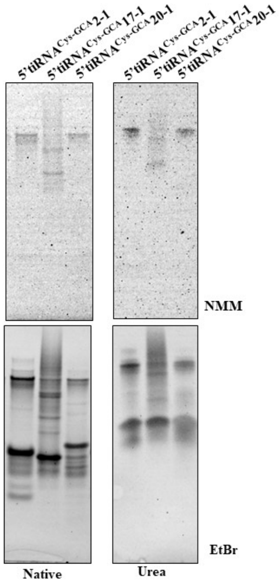

**Supplementary Figure Legends**

***Supplementary Figure S1*. (A)** Multi-sequence alignment of the 5'-tiRNAs used for *in vitro* translation assay. (**B)** Concentration-dependent translation repression of capped *Nano luciferase* reporter mRNA in HEK293-based translation extract by 5'-tiRNA$^{His\text{-}GTG}$ and 5'-tiRNA$^{Gly\text{-}GCC}$.

***Supplementary Figure S2***. (**A)** Multi-sequence alignment of various isodecoders of tRNA$^{Cys\text{-}GCA}$ using MultAlin software tool. Amongst the 29 isodecoders of tRNA$^{Cys\text{-}GCA}$ present in humans, all isodecoders possess five guanosines at the 5'-end except tRNA$^{Cys\text{-}GCA}$ (17-1, 20-1 and 22-1). (**B**) Secondary structure prediction (using RNAFold software) of different isodecoders of 5'-tiRNA$^{Cys\text{-}GCA}$ used for *in vitro* translation assays. (**C**) *In vitro* translation assays of capped and uncapped *Firefly luciferase* reporter mRNAs in HEK293 translation extracts in the presence of different isodecoders of 5'-tiRNA$^{Cys\text{-}GCA}$. (**D**) Different isodecoders of 5'-tiRNA$^{Cys}$ run in a 20% TBE native gel or 15% urea gel after folding in KCl Buffer supporting G-quadruplex assembly. The gel was first stained with 1μm NMM (dye specific for parallel G4s) followed by destaining and staining with EtBr dye (general staining).
